# Floral vibrations by buzz-pollinating bees achieve higher frequency, velocity and acceleration than flight and defence vibrations

**DOI:** 10.1101/2019.12.17.879981

**Authors:** David J. Pritchard, Mario Vallejo-Marín

## Abstract

Vibrations play an important role in insect behaviour. In bees, vibrations are used in a variety of contexts including communication, as a warning signal to deter predators and during pollen foraging. However, little is known about how the biomechanical properties of bee vibrations vary across multiple behaviours within a species. In this study, we compared the properties of vibrations produced by *Bombus terrestris audax* (Hymenoptera: Apidae) workers in three contexts: during flight, during defensive buzzing, and in floral vibrations produced during pollen foraging on two buzz-pollinated plants (*Solanum*, Solanaceae). Using laser vibrometry, we were able to obtain contactless measures of both the frequency and amplitude of the thoracic vibrations of bees across the three behaviours. Despite all three types of vibrations being produced by the same power flight muscles, we found clear differences in the mechanical properties of the vibrations produced in different contexts. Both floral and defensive buzzes had higher frequency and amplitude velocity, acceleration, and displacement than the vibrations produced during flight. Floral vibrations had the highest frequency, amplitude velocity and acceleration of all the behaviours studied. Vibration amplitude, and in particular acceleration, of floral vibrations has been suggested as the key property for removing pollen from buzz-pollinated anthers. By increasing frequency and amplitude velocity and acceleration of their vibrations during vibratory pollen collection, foraging bees may be able to maximise pollen removal from flowers, although their foraging decisions are likely to be influenced by the presumably high cost of producing floral vibrations.

## Introduction

Vibrations play an essential role in the natural behaviour of animals, particularly, among invertebrates. For example, spiders and antlions use vibrations produced by prey during hunting (Guillette et al., 2009; Mencinger-Vračko & Devetak, 2008; Nakata, 2010), and larval leafminers use vibrations to detect and avoid parasitoid wasps (Djemai et al., 2001). Animal vibrations can be transmitted both through the air (sound) and through the underlying substrate (most often plant tissue) as substrate-borne vibrations (Cocroft & Rodríguez, 2005). The substrate-borne component of vibrations can be particularly important in some contexts such as during insect communication because vibrations produced by small animals can be more efficiently transmitted through the substrate than through air (i.e. as sound) (Barth et al., 2005; Cocroft and Rodríguez, 2005; Mortimer, 2017).

Most studies of insect vibrations have focussed on vibrations produced for communication or as a by-product of flight (Hill et al., 2019; Tercel et al., 2018). But insects can use vibrations for much more than communication and locomotion. Among bees, vibrations play a particularly multifaceted role. For example, bees not only use vibrations to communicate with their nest mates (Barth et al., 2005) and as a warning or defence mechanism against potential predators (Hrncir et al. 2008; Barth et al., 2005), but also during nest construction (Rosenheim, 1987), and as a foraging tool to harvest pollen from certain flowers (Macior, 1962; Thorp, 2000; Vallejo-Marín, 2019). Substrate-borne vibrations are also the mechanism by which some bees dislodge and collect pollen on flowers with poricidal anthers (anthers that release pollen through small pores or slits; (Buchmann, 1983). The ability to use vibrations during pollen harvesting occurs in approximately 58% of all bee (Anthophila) species including 15% of genera in all bee families (Cardinal et al., 2018), and buzz-pollination (pollination using vibrations) is associated with more than 20,000 species of flowering plants (Buchmann, 1983; De Luca & Vallejo-Marín, 2013). Despite the widespread use of vibrations across diverse behavioural contexts, including during buzz pollination, we still know relatively little about the extent to which vibrational properties vary within the same species and across behaviours.

In bees, the same mechanism that drives the wings during flight is responsible of producing vibrations used during communication, defence and buzz pollination. Vibrations are produced by cyclical deformations of the bee’s thorax caused by the alternate contraction of dorsal longitudinal and dorso-ventral power flight muscles (Hedenström, 2014). These contractions are not synchronised with nerve impulses, instead bee flight muscles are “stretch-activated”, with the stretching of one of the antagonistic pairs of muscles stimulating the contraction of the other. This cycle of stretching and contraction creates a relatively self-sustaining series of cyclical thorax contractions along longitudinal and ventral axes (Dickinson, 2006; Josephson et al., 2000), with nerve impulses mostly working to maintain this cycle or make broad-scale changes such as an increase in power (Gordon & Dickinson, 2006).

Despite sharing a common production mechanism (thoracic power flight muscles), flight and non-flight vibrations in bees have clearly different vibrational properties. Non-flight vibrations are produced with the wings folded, effectively uncoupling power flight muscle contraction and wingbeat (King et al., 1996). For a given bee species, non-flight vibrations have higher frequencies than those produced during flight (Barth et al., 2005; De Luca et al., 2019; Hrncir et al., 2008; King & Buchmann, 2003), in part due to reduced drag from the wings as well as increased tension in the thoracic muscles (Hrncir et al., 2008; King et al., 1996). In contrast, non-flight vibrations produced in different contexts are superficially very similar. Both defence and floral vibrations are produced with folded wings and it is not clear to what extent non-flight thoracic vibrations have different properties to one another. Few studies have compared non-flight vibrations produced in different contexts on the same bee species. Hrncir et al. (2008) found that the frequency of vibrations produced by the tropical stingless bee, *Melipona quadrifasciata* Le Peletier (1836) (Apidae), during defence buzzes is approximately 60% of the frequency of vibrations used to communicate between foragers (350 vs. 487 Hz, respectively). In bumblebees (*Bombus spp.* Lattreille 1802), comparison of two European species found frequency differences in non-flight vibrations, namely defence and floral buzzes. However, the direction and size of the difference in frequency between defence and floral buzzes differed between the two bumblebee species (De Luca et al., 2014). While non-flight vibrations in bees are a potentially useful system for understanding the evolution and diversification of vibratory behaviours, clearly, more work is needed to characterise the exact differences between non-flight vibrations in different contexts.

Comparing the properties of vibrations produced on different behavioural contexts is technically challenging. Traditionally, substrate-borne vibrations produced by bees have been studied indirectly by recording the airborne component of the vibration using acoustic recorders. Yet, recent work indicates that although frequency components are reliably inferred from either acoustic or substrate-borne measurements, the magnitude of substrate-borne vibrations are poorly correlated with the magnitude of their acoustic component (De Luca et al., 2018). This may be because small invertebrates are poor acoustic transducers (De Luca et al., 2018), a view that is consistent with the fact that most insect communication occurs through a plant substrate, rather than through airborne sound (Cocroft & Rodríguez, 2005). This is one reason why most of the previous work comparing the vibration properties of different bee behaviours has been focused on acoustically measured frequency differences, with relatively few studies attempting to measure both frequency and amplitude (acceleration, velocity or displacement) components (Hrncir et al., 2008). To get a complete view of how vibrations differ across bee behaviours, it is necessary to capture both frequencies and amplitudes components (Vallejo-Marín, 2019). Vibration amplitude can be experimentally measured using vibration transducers such as accelerometers or laser vibrometers (Cocroft & Rodríguez, 2005). A full characterisation of substrate-borne vibrations is particularly important in the context of buzz pollination because biophysical models of poricidal anthers (Buchmann & Hurley, 1978), as well experimental tests with artificial buzzes, suggest that vibration amplitude, rather than frequency, is a key determinant of the rate of pollen ejection from flowers (De Luca et al., 2013; Rosi-Denadai et al., 2018).

In this study, we use accelerometers and laser vibrometry to measure the vibrational properties of buzzes produced by bumblebees (*Bombus terrestris ssp. audax*, (Harris 1776); hereafter *B. audax*) in three different behavioural contexts: Flight, defence and floral vibrations. In addition, we compare the floral vibrations produced by bees on two different buzz-pollinated plant species (*Solanum rostratum* Dunal and *S. citrullifolium* (A. Braun) Nieuwl., Section *Androceras*, Solanaceae). Previous work has shown conflicting results on the extent to which bumblebees change the vibrations produced during floral visitation (floral vibrations), with some studies showing differences between flowers (Switzer and Combes, 2017) or with experience (Morgan et al., 2016; Switzer et al., 2019) and others showing more limited flexibility (Russell et al., 2016b). However, to date, few studies have used non-contact methods (laser vibrometry) to look specifically at the vibration produced by the bee rather than those transmitted through different flowers (Nunes-Silva et al., 2013). Our study addresses three specific questions: 1) What are the main differences in the vibrations produced by bumblebees across different behaviours? 2) To what extent floral vibrations produced by the bee depends on the species of flower being visited? 3) Do the characteristics of vibrations depend on bees’ morphological traits such as size?

## Methods

### Study system

#### Bees

We used two colonies of the buff-tailed bumblebee, *Bombus terrestris audax* (Koppert, Agralan Ltd, Wiltshire, UK). Each colony had access to *ad libitum* “nectar” solution (Koppert) within the colony. Each colony was attached to a flight arena (122 × 100 × 37 cm), illuminated with an LED light panel (59.5 × 59.5 cm, 48 W Daylight; Opus Lighting Technology, Birmingham, UK) and maintained on a 12h:12h supplemental light:dark cycle. The ambient temperature was 20-23°C and humidity was 50-60% RH. In each arena, bees were also provided with a 1M sucrose solution, *ad libitum*, from three feeders in each colony, as well as eight inflorescences (four *Solanum rostratum*, four *S. citrullifolium*) every two days.

#### Plants

We tested floral vibrations on two closely related species from the genus *Solanum* (Solanaceae). *Solanum rostratum* and *Solanum citrullifolium* are both nectarless species, which attract and reward pollinators solely with pollen. In common with other *Solanum* species, *S. rostratum* and *S. citrullifolium* have poricidal anthers, which requires pollinators to vibrate the anthers to release pollen. Unlike some other *Solanum* species, *S. rostratum* and *S. citrullifolium* are both heterantherous, with bees primarily focussing their attention on “feeding anthers” presented at the centre of the flower, while a single, rarely visited “pollination anther” deposits pollen on the visiting bee. *Solanum* species are a classic system for the study of buzz pollination (e.g. Buchmann & Cane, 1989; King & Buchmann, 1996), and *S. rostratum* and *S. citrullifolium* have been directly compared in a previous study which identified apparent difference in the coupling factors of these species (Arroyo-Correa et al., 2019), making this pair an ideal comparison.

*S. rostratum* and *S. citrullifolium* plants were grown from seed at the University of Stirling research glasshouses, using the method described in Vallejo-Marín et al. (2014). Seeds of *S. rostratum* were collected in Mexico (20.901°N, 100.705°W; accessions 10s77, 10s81, 10s82) and seeds of *S. citrullifolium* were obtained from self-fertilised fruits (accession 199) grown from seeds obtained from Radboud University’s seed collection (accession 894750197). For daily flower provision for bees, inflorescences were placed in water-soaked Ideal Floral Foam (Oasis Floral Products, Washington, UK) in plastic containers. For experiments, we used a single flower, cut 2-3cm below the calyx.

#### Recording of floral vibrations

To facilitate the recording of bee vibrations using laser vibrometry, we tagged individual bees with a small (2mm^2^) piece of reflective tape placed in the dorsal part of the thorax. Bees buzzing on flowers in the flight cages were captured, placed in a freezer at −26°C for seven minutes, and tagged with reflective tape using Loctite UltraControl instant adhesive (Henkel Limited, Winsford, UK). After being at room temperature, bees resumed normal activity after approximately 7-10 minutes and were released back into the colony.

At least 24 hours after being tagged, bees were allowed to visit flowers in the arena and a tagged bee which was actively buzzing flower was collected from flowers in the flight cage and released onto a single flower of either *S. rostratum* or *S. citrullifolium* in the test arena. The flower species were chosen so that each colony received the same number of flowers from each plant species. The vibrations produced by the bee were recorded simultaneously in two ways. First, we measured vibrations produced in the bee’s thorax using a laser vibrometer (PDV 100, Polytec, Coventry, UK). Laser vibrometry provides a direct, contactless measure of the vibrations produced by the bee. Vibrations measured with the laser were sampled at a rate of 10240 Hz using a low pass filter of 5Hz, and a maximum velocity range of either 100 mm/s (for bees 1-14) or 500 mm/s (for bees 15-32). The laser vibrometer was placed approximately 20cm away from the flower and aimed at the reflective tag on the bee’s thorax. Second, we used an accelerometer (352C23, 0.2g; PCB Piezotronics) to record the vibrations transmitted from the bee to the flower (Arroyo-Correa et al., 2018). The accelerometer was attached to the calyx at the base of the flower being vibrated by the bee using a 5mm × 0.35mm pin made from an entomological pin (Austerlitz, Size 0) and glued to the accelerometer with instant adhesive as described in Arroyo-Correa et al. (2018). The accelerometer and laser were set to register along the same axis of movement.

Both laser vibrometer and accelerometer data were simultaneously recorded and time-stamped using Data Acquisition System (cRIO model 9040 with the C series module NI 9250; National Instruments, Newbury, UK) using a custom-made LabView (National Instruments) program (available upon request). While the bee buzzed the flower, data were recorded during two seconds at a sampling rate of 10240 Hz and saved to a file. After collecting 5-10 buzzes for each bee, the bee was caught in a 30mL plastic container (201150; Greiner, Gloucestershire, UK), and euthanised by being placed in −26 freezer for 48 hours. In total, we collected data for 16 bees from two colonies, eight on each flower species. For each bee we recorded analysed an average of 6.13 buzzes (N = 98 buzzes from 16 bees).

#### Recording of defence and flight vibrations

For the recording of flight and defence buzzes bees were selected at random from the flight box. As for the flower buzzing, bees were immobilised by being placed in the freezer for seven minutes. In addition to gluing a 2mm^2^ reflective tag to the scutum, immobile bees were also tethered to the apparatus for recording defence and flight buzzes, similar to that used by Hrncir et al. (2008). The neck of the bee was held by a loop of fine nylon string threaded through a needle and attached to a syringe secured by a clamp (Figure 1). After 7-10 minutes, the tethered bee had returned to regular activity levels and we continued with data collection.

**Figure 1:**
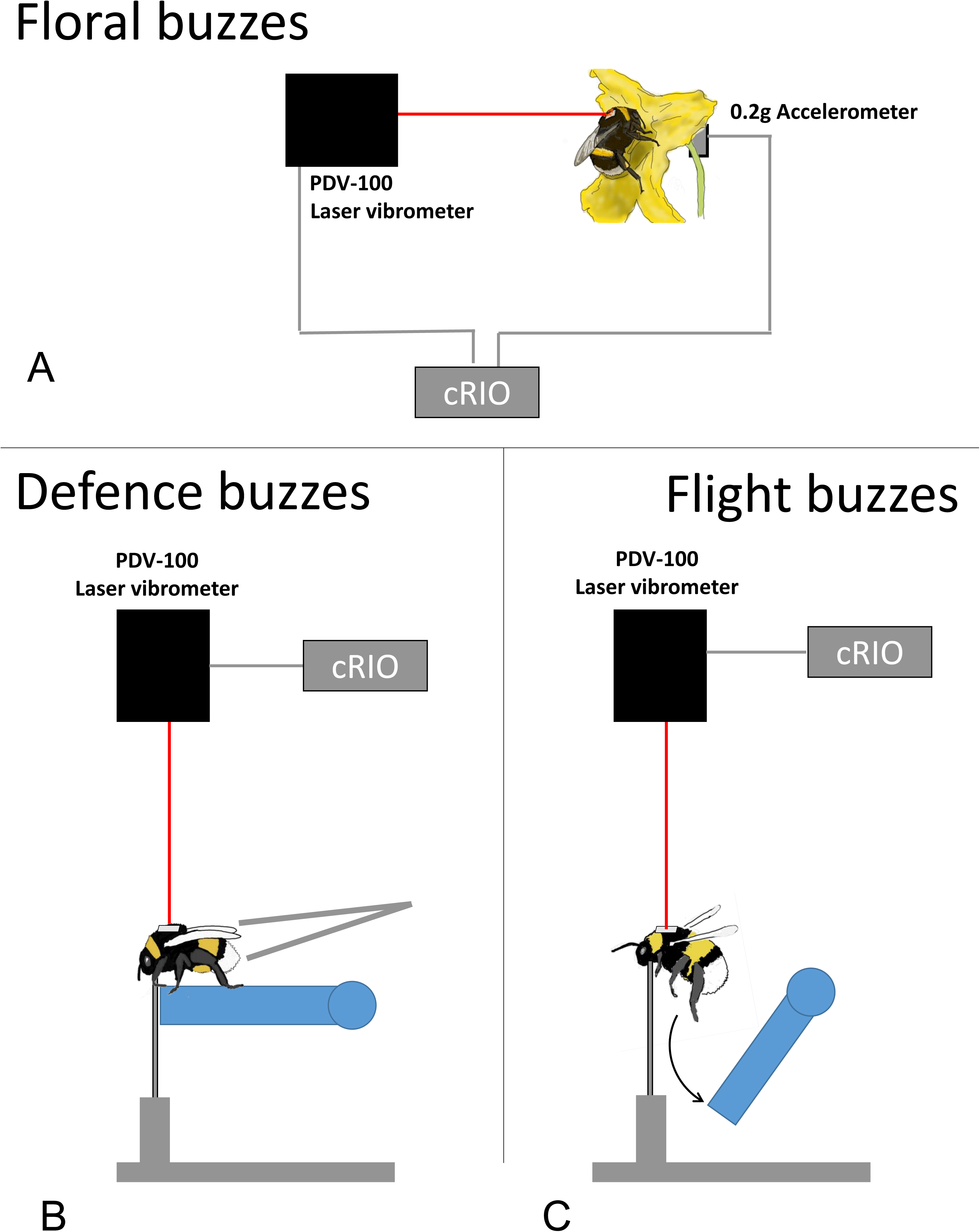
Experimental set up for measuring bee vibrations. For floral buzzes (top), vibrations were recorded simultaneously by a PDV-100 laser vibrometer focussed on a 2mm^2^ reflective tag on the back of the thorax of the bee, and by a 0.2g accelerometer pinned to the calyx at the base of the flower. These measurements were sent to the compactRIO data acquisition unit (cRIO) which timestamped the data and exported them to a file. For defence and flight buzzes (bottom), bees were tethered on a platform using a nylon wire loop fed through a blunted needle. For defence buzzes (left), bees were gently squeezed on the abdomen using featherweight tweezers. To stimulate flight (right), the platform was dropped away triggering reflexive flight. In both cases, vibrations were recorded using a PDV-100 laser vibrometer positioned above the bee and aimed at a 2mm^2^ on the back of the thorax. The vibrometer then send the data to the cRIO to be timestamped and exported.

To record both flight and defence buzzes, the laser vibrometer was placed above the bee and aimed at the tag on the bee’s thorax. The laser beam was perpendicular to the platform on which the bee was tethered. Defence and flight vibrations measured with the laser were sampled at a rate of 10240 Hz using a low pass filter of 5Hz, and a maximum velocity range of 500 mm/s. To induce defence buzzes, the tethered bees were gently squeezed along the sides using featherweight forceps. To record flight buzzes, the platform underneath the tethered bee fell away inducing the bee to start flight activity (Hrncir et al., 2008). As before, vibration data was recorded through the cRIO data acquisition system using a custom LabVIEW program, which collected two seconds of data at a time at a sampling rate of 10240 Hz, with a low pass filter of 5Hz and a velocity range of 500 mm/s. Flight and defence buzzes were recorded from 20 bees in total, with defence and flight buzzes captured from all bees. To avoid order effects, 10 of the bees had defence buzzes collected first and 10 had flight collected first. Following recording, tethered bees were immobilised again by being placed in the freezer, removed from the tether, placed in a plastic container, and euthanised in the − 26°C freezer. For each bee, we analysed an average of 5.6 flight vibrations (n = 112 vibrations from 20 bees) and 6.8 defence buzzes (n = 136 from 20 bees).

### Bee size

Bee size was approximated using intertegular distance (ITD), the distance between the tegulae at the base of the wings (Cane, 1987). We measured ITD using a digital photograph of euthanised bees taken with a dissecting microscope (MZ6, Leica Microsystems, Milton Keynes, UK) (Figure S1), and analysed with the *FIJI* distribution of *ImageJ* (Schindelin et al., 2012).

### Data Analysis

#### Analysing vibrations

We used a section of each recorded vibration for analysis (Figure 2). For floral buzzes, we selected a section of each recording that successfully captured both laser and accelerometer sensors. The sensor data (time series with voltage units) were converted from voltage to either velocity (laser) or acceleration (accelerometer) using the factory-provided conversion factors for each sensor. We zero-centred the data by subtracting the mean amplitude from each value and applied an 80-5000 Hz band-pass filter and a Hamming window (window length = 512), using the *fir* function in the *R* package *seewave* (Sueur et al., 2008). The acceleration data were converted to velocity by numerical integration using the *cumtrapz* function in the *pracma* package (Borchers, 2019), and the band-pass filter was applied again. The fundamental frequency of the analysed vibration was obtained with the *fund* function, calculated over the entire sample and setting a maximum frequency to 1000 Hz. Peak amplitude velocity for each vibration segment was calculated from the amplitude envelope calculated using the *env* function with a mean sliding window of length 2 and an overlap of 75%. All analyses were done in *R* version 3.6.0 (R Core Team, 2019)

**Figure 2.**
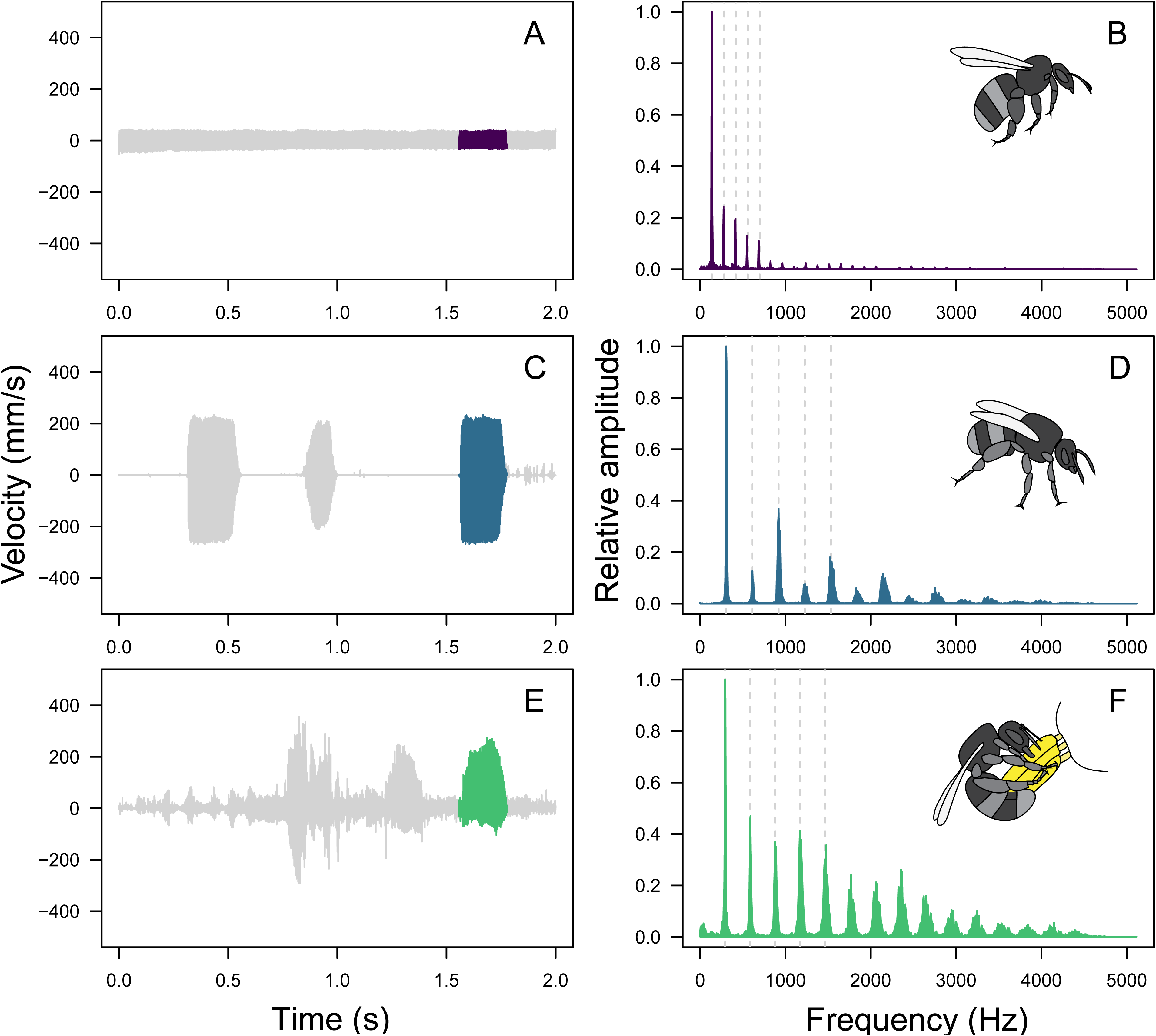
Oscillograms and frequency spectra of vibrations (buzzes) produced by bumblebees (*Bombus terrestris audax*) in three different behavioural contexts: Flight (A, B), defence (C, D), and buzz pollination (E, F). Left-hand side panels (A, C, E) show buzzes in the time domain (oscillograms), while right-hand side panels show buzzes in the frequency domain (frequency spectra; B, D, F). The coloured region in the oscillogram show the section of the buzz used to generate the corresponding frequency spectrum. The first five harmonics (multiples of the fundamental frequency) are shown as vertical dashed lines in the frequency spectra.

#### Transmission of bee vibrations through flowers

To quantify the extent to which the vibrations produced by bees differ from those measured in the flower itself, we calculated King’s coupling factor (King, 1993). The bee’s coupling factor (*Kbee*) was calculated by dividing the root mean squared (RMS) amplitude velocity of the vibration produced by the bee by the RMS amplitude velocity recorded by the accelerometer placed in the flower’s calyx (Arroyo-Correa et al. 2019). We also calculated King’s coupling for vibrations produced by a mechanical calibrated shaker (Handheld shaker model 394C06, PCB Piezotronics). The calibrated shaker produces a vibration of constant properties (frequency = 159.2Hz, RMS amplitude velocity = 9.8 mm s^−1^) that are transmitted to a small metal plate at one end of the instrument. The metal plate of the calibrated shaker was firmly pushed against the feeding anthers of the flower, and we recorded four to five samples of two seconds each using the data acquisition system described above (*Analysing Vibrations*). For each flower, we selected one clean recording, converted voltage to velocity as described above, and obtained King’s coupling factor for the shaker (*Kshaker*) using the ratio between expected and observed RMS velocity. Measuring both *Kbee* and *Kshaker* allowed us to compare the difference in the efficiency with which a bee and a mechanical shaker transmit vibrations to the flower.

#### Statistical analyses

To compare the properties of vibrations in different contexts we used linear mixed effect models using either peak velocity or fundamental frequency as response variables, buzz type (flight/defence/floral) and intertegular distance as explanatory variables, and bee identity as a random effect. In addition to peak velocity and frequency, which were measured directly, we also used these measures to derive the displacement amplitude (in mm) and acceleration (in mm/s^2^) of the vibration. As with velocity, we analysed the peak recordings of each of these measures with linear mixed effect models, with buzz type and intertegular distance as explanatory variables and bee identity as a random effect. To compare the properties of floral vibrations on different *Solanum* species, we employed linear mixed effect models, using either laser-recorded peak velocity, laser-recorded fundamental frequency, accelerometer-recorded peak velocity or accelerometer-recorded fundamental frequency as response variables, flower species and intertegular distance as explanatory variables, and bee identity as a random effect. Finally, to compare the effect of flower species and recording method on coupling factors, we used a linear mixed effect model with coupling factor as a response variable, flower species, intertegular distance, and vibration method (bee vs artificial) as explanatory variables, and bee ID as a random effect. All analyses were performed using *lme4* (Bates et al., 2015) to estimate parameters and *lmerTest* (Kuznetsova et al., 2017) to assess statistical significance.

## Results

### Comparison of buzzes produced in different behavioural contexts

The vibrations produced during flight, defence and pollen extraction differ significantly in properties including fundamental frequency and peak amplitude velocity (Table 1). The peak amplitude velocity of floral buzzes (262.85 ± 9.52 mm/s) was significantly higher than both defence (194.85 ± 6.12 mm/s) and flight buzzes (57.29 ± 1.28 mm/s; Figure 3A, Table 1). We found no significant effect of bee size on peak amplitude velocity (Table 1). Floral buzzes also had significantly higher frequencies (313.09 ± 2.63 Hz) than both defence (236.32 ± 4.29 Hz) and flight buzzes (136.95 ± 1.73 Hz) (Figure 3B). We also detected an interaction between bee size and buzz type with larger bees achieving higher frequency defence buzzes and lower frequency flower and flight buzzes than smaller bees (Table 2). The differences in peak amplitude velocity across the three behaviours observed here extended to peak amplitude acceleration, with floral buzzes achieving higher accelerations (517.77m s^−2^ ± 19.40), than defence (297.41m s^−2^ ± 11.96), and flight vibrations (49.43 m s^−2^ ± 1.34) (Figure 3D). In contrast, the peak amplitude displacement of floral (0.27 mm ± 0.009) and defence buzzes (0.27 mm ± 0.007) were similar, although both greater than the displacement amplitude of flight vibrations (0.14 mm ± 0.005) (Figure 3C).

**Figure 3.**
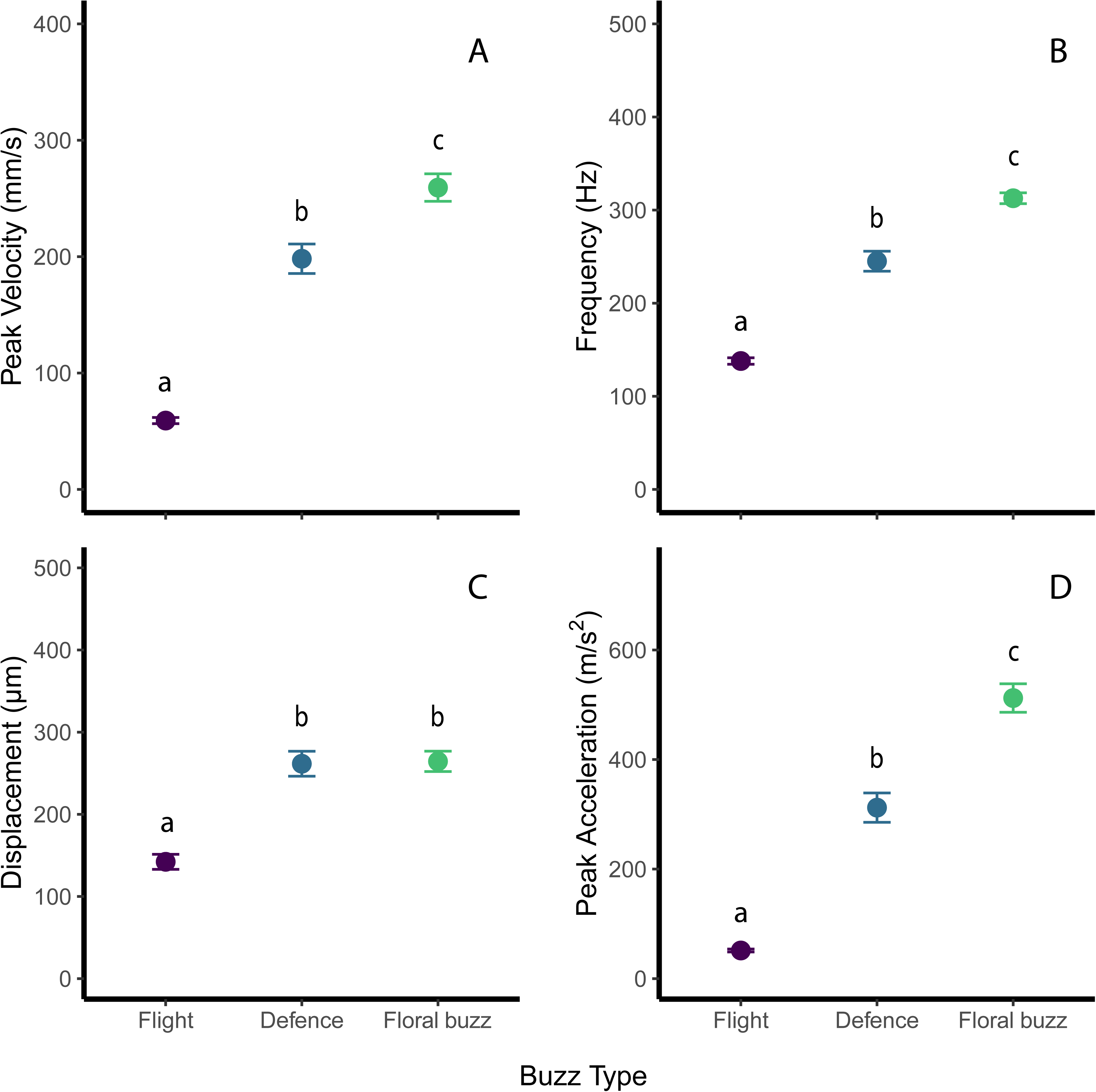
Differences in the properties of vibrations (buzzes) produced in different contexts (flight, defence, floral buzzes). Vibrations differed in both peak velocity (A) and frequency (A), with floral buzzes exhibiting the highest velocity and highest frequency buzzes, and flight producing the lowest velocity and frequency vibrations. From these values we derived the magnitude of the vibrations, in terms of displacement of the thorax, (C) and the acceleration (D) produced during these vibrations. Although there was no difference in the absolute magnitude of the vibrations produced during defence and floral buzzes, because the floral buzzes were faster and at higher frequency than the defence buzzes, floral buzzes showed much higher accelerations.

**Table 1.** Analysis of bee size (intertegular distance) and behavioural context on the properties of thoracic vibrations measured with a laser vibrometer. The parameter estimates and standard errors were calculated from a linear mixed effect model with bee identity as a random factor. *P*-values for each explanatory variable were calculated using a Type III analysis of variance with Satterthwaite’s estimation of degrees of freedom. Statistically significant values are in bold.

| Response variable | Parameter | Estimate | SE | *P*-value |
| --- | --- | --- | --- | --- |
| **Peak Amplitude Velocity (mm/s)** | Intercept (Buzz Type: Flight) | 165.71 | 94.16 |  |
|  | Intertegular distance | -24.63 | 21.72 | 0.27 |
|  | Buzz Type |  |  | **< 0.001** |
|  | Defence | 132.68 | 8.54 |  |
|  | Floral | 207.65 | 14.53 |  |
| **Fundamental Frequency (Hz)** | Intercept (Buzz Type: Flight) | 200.93 | 70.89 |  |
|  | Intertegular distance | -14.53 | 16.36 | 0.38 |
|  | Buzz Type |  |  | **< 0.001** |
|  | Defence | 102.93 | 3.38 |  |
|  | Floral | 177.70 | 10.50 |  |
|  | Buzz Type*Intertegular distance |  |  | **0.002** |
| **Displacement (mm)** | Intercept (Buzz Type: Flight) | 0.24 | 0.11 |  |
|  | Intertegular distance | -0.022 | 0.026 | 0.40 |
|  | Buzz Type |  |  | **< 0.001** |
|  | Defence | 0.11 | 0.011 |  |
|  | Floral | 0.13 | 0.017 |  |
| **Acceleration (m/s<sup>2</sup>)** | Intercept (Buzz Type: Flight) | 358.32 | 199.45 |  |
|  | Intertegular distance | -71.09 | 46.01 | 0.13 |
|  | Buzz Type |  |  | **< 0.001** |
|  | Defence | 248.57 | 16.82 |  |
|  | Floral | 479.57 | 30.57 |  |

**Table 2.** Analysis of bee size (intertegular distance), plant species, and recording location on the properties of floral vibrations. Vibrations were recorded on *S. citrullifolium* and *S. rostratum*, both directly on the bee’s thorax using a laser vibrometer and on the flower using an accelerometer. The parameter estimates and standard errors were calculated from a linear mixed effect model with bee identity as a random factor. *P*-values for each explanatory variable were calculated using a Type III analysis of variance with Satterthwaite’s estimation of degrees of freedom. Statistically significant values are in bold.

| Response variable | Variable | Estimate | SE | *P*-value |
| --- | --- | --- | --- | --- |
| **Peak Amplitude Velocity (mm/s)** | Intercept (Plant: *S. citrullifolium*, Location: Bee) | 312.06 | 74.43 |  |
|  | Intertegular distance | -13.74 | 16.31 | 0.42 |
|  | Plant species: *S. rostratum* | 22.22 | 12.95 | 0.11 |
|  | **Location: Flower** | **-233.35** | **9.30** | **<0.001** |
| **Fundamental Frequency (Hz)** | Intercept (Plant: *S. citrullifolium*, Location: Bee) | 462.66 | 60.83 |  |
|  | **Intertegular distance** | **-33.54** | **13.36** | **0.027** |
|  | Plant species: *S. rostratum* | 4.40 | 10.12 | 0.67 |
|  | Location: Flower | -0.20 | 2.07 | 0.92 |

### Floral buzzes

Our analyses of the vibrations produced by bees while visiting flowers (floral buzzes) shows that only some of the properties of these vibrations depend on whether they are recorded on the bee or on the flower (Figure 4). The magnitude of vibrations recorded directly on the bee had considerably higher peak velocity amplitudes (273.56 ± 12.49 and 247.34 ± 14.53 mm/s for *S. rostratum* and *S. citrullifolium* respectively) than those vibrations measured on the flower (36.61 ± 2.30 and 19.20 ± 1.03 mm/s for *S. rostratum* and *S. citrullifolium*, respectively; Figure 5A, Table 2). In contrast, the fundamental frequency of the floral vibrations was similar whether recorded directly from the bee (313.16 Hz ± 2.86 and 312.09 Hz ± 4.99 Hz for *S. rostratum* and *S. citrullifolium*, respectively) or indirectly via the accelerometer on the flower (312.70 Hz ± 2.92 and 313.16 Hz ± 4.81 for *S. rostratum* and *S. citrullifolium*, respectively; Figure 5B, Table 2). Interestingly, we observed that vibrations measured on the bee contained more harmonics (*S. citrullifolium*: 10.75 ± 0.38; *S. rostratum*: 11.34 ± 0.35) than those observed on vibrations measured on the flower (*S. citrullifolium*: 3.65 ± 0.27; *S. rostratum*: 2.57 ± 0.20) (Figure 4).

**Figure 4.**
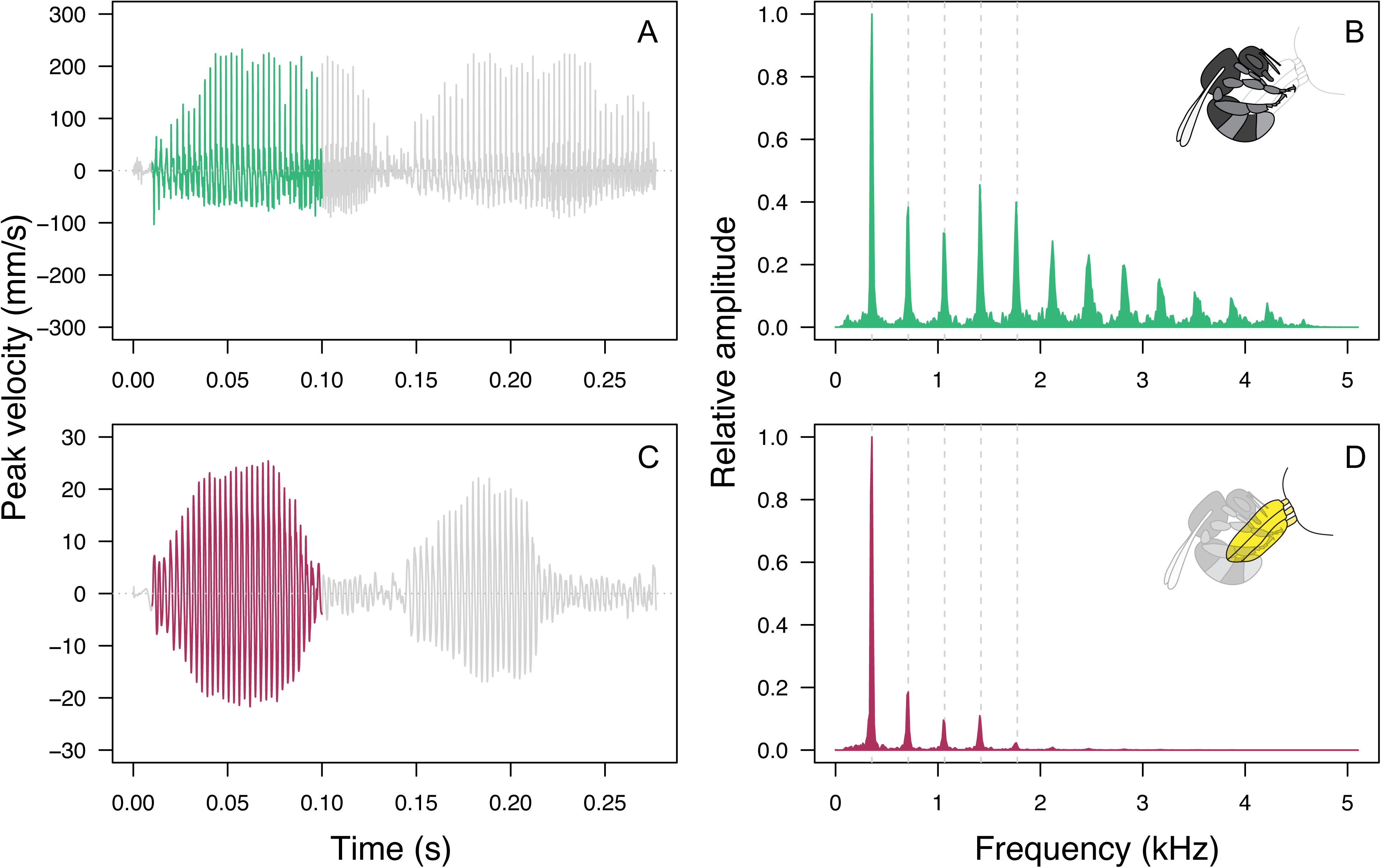
Example of a floral vibration produced by *Bombus terrestris audax* while visiting a flower of *Solanum citrullifolium* as recorded directly from the bee (A, B) and on the flower (C, D). The magnitude of the vibration, measured as peak velocity amplitude, is much higher when measured directly on the bee’s thorax with a laser vibrometer (A), than when measured using an accelerometer attached to the base of the flower (C). In contrast, the fundamental frequency of the buzz produced during floral visitation is the same (355 Hz) whether is measured in the bee’s thorax (B) or on the base of the flower (D). The coloured section in the oscillograms shown in A and C represent the section of the buzz used to calculate the frequency spectra shown in B and D. The dashed lines in panels B and D represent the first five harmonics of the fundamental frequency.

**Figure 5.**
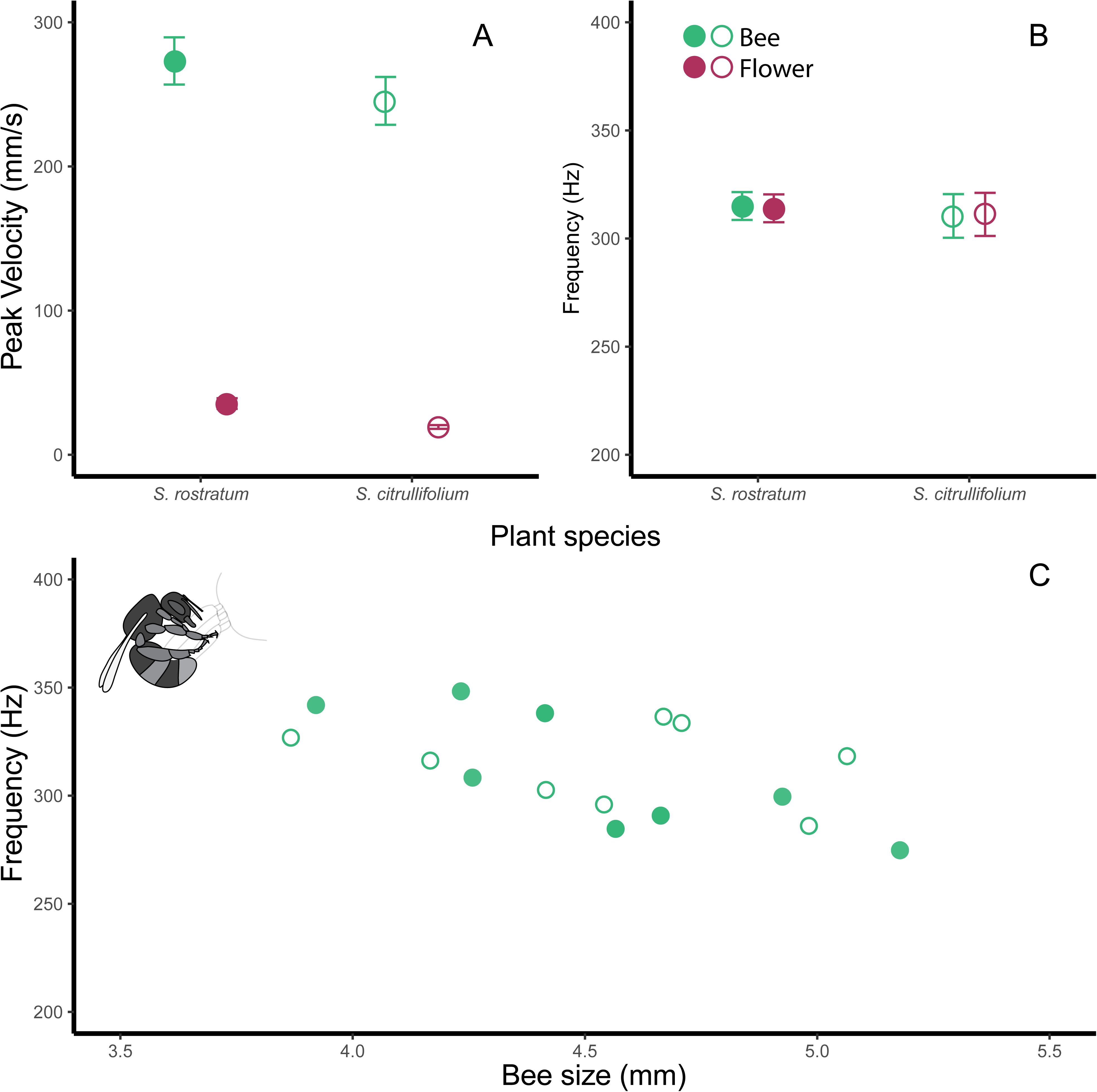
Peak amplitude velocity (A) and fundamental frequency (B) of floral buzzes of *Bombus terrestris audax* on buzz-pollinated flowers of *Solanum rostratum* (closed symbols) and S*. citrullifolium* (open symbols). Floral buzzes were recorded directly from the bee‘s thorax using a laser vibrometer (green symbols) or on the flower using an accelerometer attached to the calyx (magenta symbols). Vibrations recorded on the flower had significantly lower peak velocities but similar fundamental frequencies as those measured in the bee. (C) Relationship between bee size (intertegular distance) and the fundamental frequency of floral buzzes. Each symbol in (C) represents the average frequency from multiple buzzes produced by an individual bee.

Plant species did not significantly affect the frequency or peak amplitude velocity of floral vibrations (but see section *Transmission of vibrations through flowers* for differences in the transmission of vibrations from bee to flower in the two *Solanum* species). Bee size (intertegular distance) was negatively associated with fundamental frequency of floral vibrations (Figure 5C), while bee size had no effect on their peak amplitude velocity (Table 2). We found no statistically significant interaction between bee size and plant species on either frequency or peak amplitude velocity of floral vibrations.

### Transmission of vibrations through flowers

To analyse the effect of plant species on the transmission of floral vibrations through the flower, we compared King’s coupling factor (*K*, the ratio of vibration magnitude produced to vibration received) for the two *Solanum* species. We found that *S. rostratum* had a significantly lower coupling factor (*Kbee* = 5.64 ± 0.61, *Kshaker* = 5.95 ± 1.77; mean ± SE) than *S. citrullifolium* (*Kbee* = 9.92 ± 0.97, *Kshaker* = 8.93 ± 1.97; Table 3, Figure 6). Our analysis showed no difference within plant species between coupling factors calculated from either bee floral buzzes (*Kbee*) or synthetic vibrations applied with the calibrated shaker (*Kshaker*) (Table 3), although *Kbee* is less variable than *Kshaker* (Figure 6). We did not find an effect of bee size on coupling factor (Table 3).

**Figure 6.**
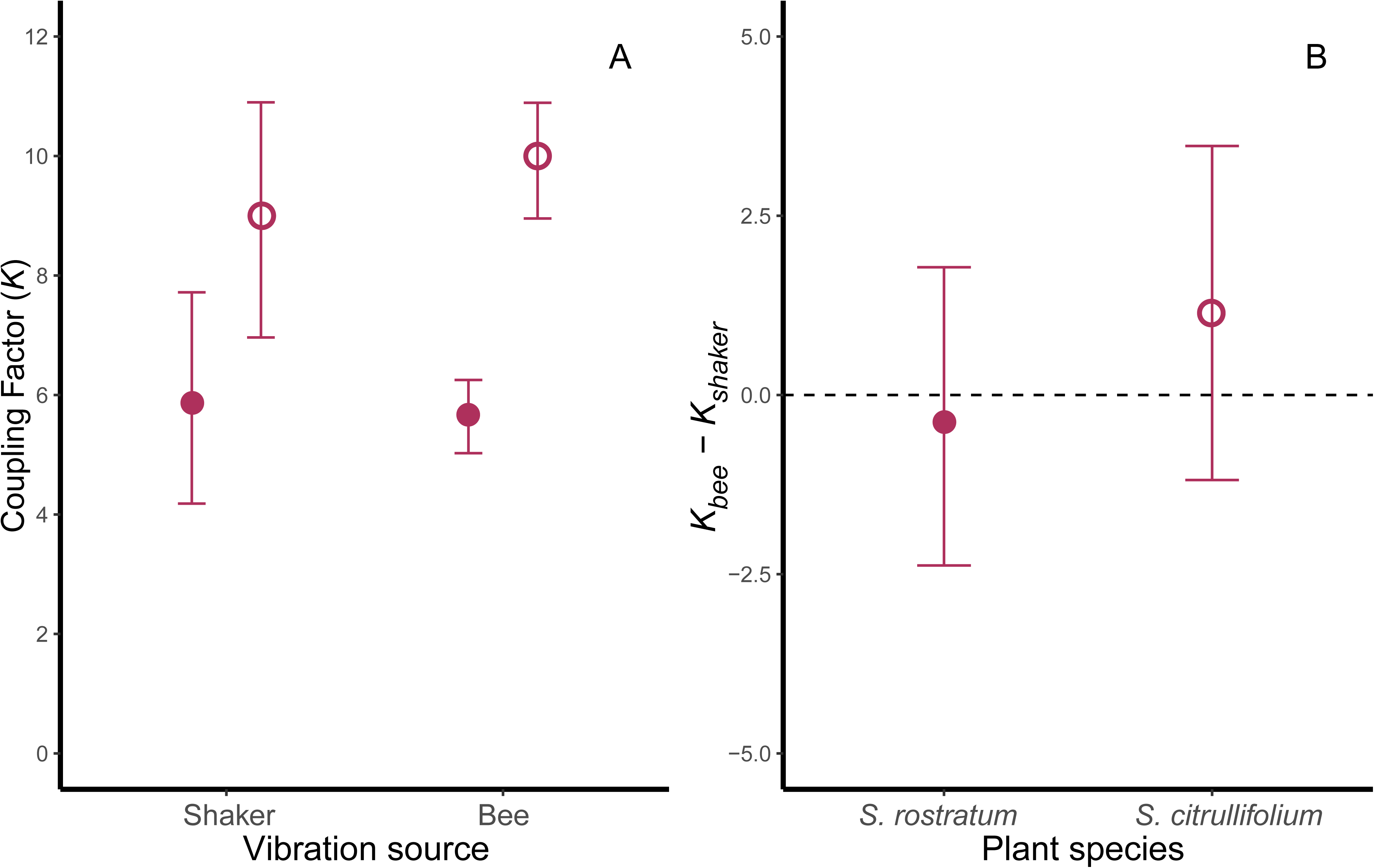
Comparison of the ratio of the magnitude of the input vibration to the magnitude of the vibration measured at the sensor (King’s coupling factor) on two buzz-pollinated species of *Solanum*. Coupling factors were estimated using either natural bee vibrations (*bee*) or synthetic vibrations produced with a calibrated mechanical shaker (*shaker*) as the input vibration. The calibrated shaker produced a vibration of fixed properties (frequency = 159.2 Hz, RMS velocity = 9.8mm/s). The magnitude of the vibration produced by the bee was measured using a laser vibrometer on the bee’s thorax. The vibration transmitted to the sensor on the flower was measured at the calyx using an accelerometer. Plant species consistently differ in their coupling factor with *S. rostratum* having lower values than *S. citrullifolium* (A), irrespective of whether it is calculated using bee or calibrated shaker vibrations (B).

**Table 3.**
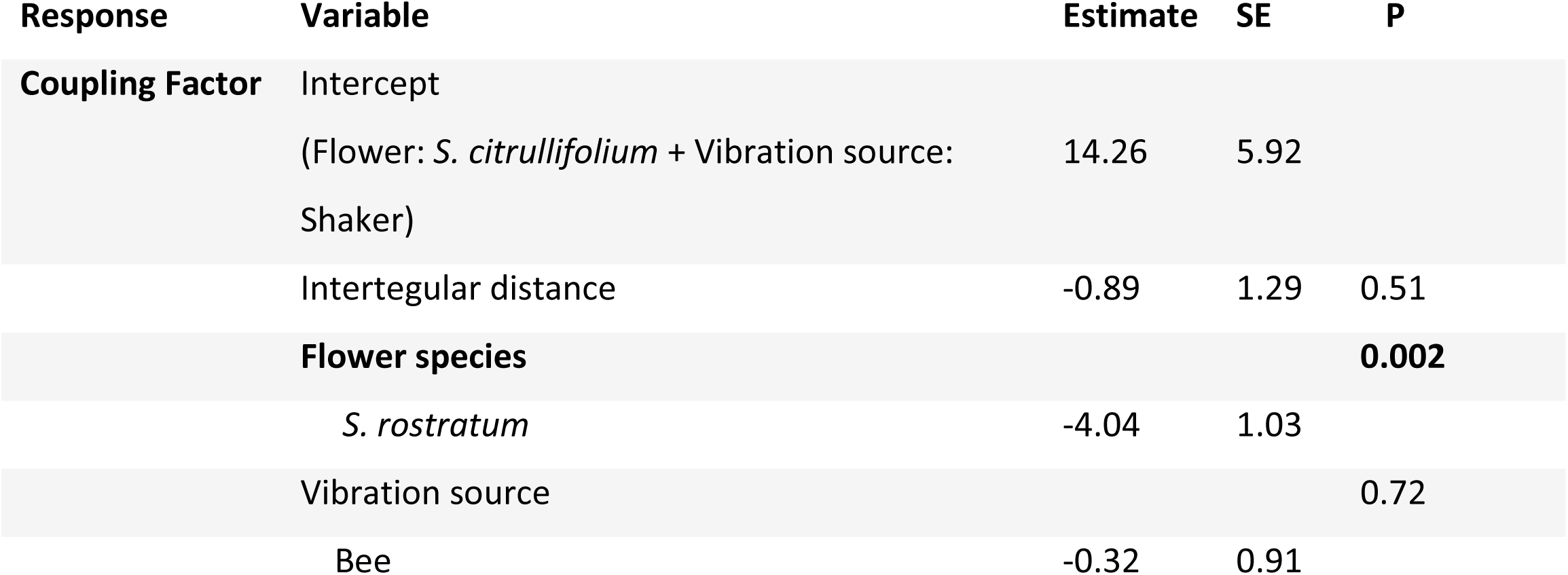
Effect of bee size (intertegular distance distance), flower species, and vibration method on the magnitude of King’s Coupling Factor. Vibrations were applied to *S. citrullifolium* and *S. rostratum*, either by the bee (bee) or by pressing a calibrated shaker against the flower (shaker). The parameter estimates and standard errors were calculated from a linear mixed effect model with bee identity as a random factor. P-values for each explanatory variable were calculated using a Type III analysis of variance with Satterthwaite’s method. Statistically significant values are in bold.

## Discussion

Bumblebees and other buzz-pollinating bees present a unique opportunity for research on insect vibrations. In addition to producing vibrations during locomotion and as a signal to predators or conspecifics, buzz-pollinating bees also use vibrations to forage. While the posture of bees during floral buzzes and defence buzzes are very similar, with both requiring the wings folded back over the body, the functions of these two buzzes are very different, making them a useful comparison for understanding how function might influence the properties of bee vibrations. Our results show clear differences in biomechanical properties of defence and floral buzzing, as well as differences between these vibrations and those produced during flight. In addition to differences between different behaviours, we also found that the species of plant being vibrated and the size of the bee, affected the properties of the floral vibrations experienced by plants.

### Floral vibrations and bee size

Our results are consistent with previous work showing that plant species differ in their transmission of floral vibrations (King 1993; Arroyo-Correa et al., 2019). Between the two studied plant species, we found that *Solanum rostratum* is better at transmitting vibrations applied on the anthers to other parts of the flower than *S. citrullifolium*, as shown by its lower coupling factor (cf. Arroyo-Correa et al., 2019). Interestingly, the coupling factor calculated using synthetic vibrations applied with a metal plate and the one calculated using vibrations applied by live bees were similar, suggesting that fine floral manipulation by the bee during buzzing has little effect on the vibrations transmitted to other parts of the flower. Further analyses of the biomechanical properties of flowers are required to determine the mechanism responsible for the different coupling factors observed here and in previous studies.

We found little evidence that the magnitude of floral, flight and defence buzzes can be explained by the range of bee size variation observed within a single species of bumblebee. In contrast, bee size was negatively associated with frequency of floral and flight buzzes but positively with defence buzzes. The frequency of flight vibrations in bees is usually negatively associated with size both within (this study) and across species (De Luca et al., 2019). For floral vibrations, the association between frequency and flight seems to vary (reviewed in De Luca et al., 2019), ranging from negative, as in our study on *B. terrestris audax*, to positive (Arroyo-Correa et al. 2019) to no detectable relationship both within species (De Luca et al., 2013; De Luca et al. 2014, Nunes et al. 2013) and across multiple species (De Luca et al., 2019; Rosi-Denadai et al., 2018). Moreover, the relationship between the frequency of floral buzzes and bee size within species may further depend on the metric of bee size used (Corbet & Huang, 2014; Switzer & Combes, 2017). Taken together this body of work suggests that differences in size are not sufficient to explain variation in floral buzzes during buzz pollination.

### Differences among buzz types

We found that bumblebees vibrating flowers produce higher accelerations than in other behaviours, and much higher than previously thought. The floral vibrations measured in this experiment were on average 500 m/s^2^, more than 2-3 times what Arroyo Correa et al. (2019) and King (1993) calculated after measuring floral buzzing from the plant and correcting with the corresponding coupling factor. Despite this, our measurements for frequency and velocity, from which acceleration was calculated, were consistent with those found by other studies looking at flying, defence buzzing, and flower buzzing bees (Nunes-Silva et al., 2003, King 1993). Floral buzzes appear to be characterised by higher accelerations, velocities, and frequencies, than defence buzzes. And both floral and defence buzzes have higher accelerations, velocities, displacement amplitude and frequencies, than are produced during flight. The key question raised by our results, then, is why are the properties of floral, defence and flight vibrations so different to one another? This question can be addressed in two ways: 1) by considering how the mechanisms underlying these vibrations might differ across behaviours; and 2) how the function of the behaviour might select for particular vibration properties.

### Mechanisms of bee vibrations

All of the vibrations we measured in this study were produced by contractions of the dorsal longitudinal and dorso-ventral flight muscles in the thorax. The fact that these vibrations all share a common mechanisms could mean that something other than the muscles might be responsible for the differences we observed. One early suggestion was whether the decoupling of the wings from the flight muscles during non-flight vibrations (defence, floral buzzes) changed the resonant properties of the thorax and led to higher frequencies. It is plausible that the deployment of the wings could lower the frequency of the vibrations, wings produce drag and inertia, which is one reason why insects with larger wing have a lower wingbeat frequency (e.g. Greenewalt, 1962; Joos et al., 1991). When insect wings are cut shorter the frequency of flight increases (Hrncir et al., 2008; Roeder, 1951). While wing deployment can explain the different between flight and non-flight vibrations, it cannot explain the differences between the two non-flight vibrations (floral and defence buzzes), where the wings remained folded and the mass of the system remains unchanged.

Instead of the mechanical effect of the wings, differences between non-flight vibrations could be the result of differences in muscle activity, either in terms of increasing muscle power or by changing the stiffness and resonant properties of the thorax. Although bumblebee flight muscles are stretch activated, and so do not contract in time with motor neuron firing, studies of similar muscles in *Drosophila* show that increasing the frequency of firing increases the Ca^2+^ concentration in the flight muscles, resulting in more powerful contractions(Dickinson et al., 1998; Gordon & Dickinson, 2006; Lehmann & Bartussek, 2017; Wang et al., 2011). Bees could also use other muscles to stiffen the thorax, changing its resonant properties, altering the frequency at which the cycle of stretch-activated contractions reaches equilibrium (Nachtigall & Wilson, 1967). Although these mechanisms have yet to be studied in bees, neurophysiological studies of bee flight muscles have found differences between flight and non-flight vibrations (Esch & Goller, 1991; King et al., 1996), which might also explain differences between non-flight vibrations. During flight, both the dorso-ventral and dorsal longitudinal muscles sets are stimulated equally, whereas during defensive buzzes the dorsal longitudinal muscles are stimulated at twice the rate as the dorso-ventral muscles (King et al. 1996). If, for example, the increased difference in activation between the flight muscles sets is responsible for the increased frequency of non-flight vibrations, then we might expect the difference in excitation between the muscle sets to be even more extreme during floral buzzes than during defence buzzes. By comparing the mechanisms underlying floral buzzes, defence buzzes, and flight, in this way, we can begin to understand how bees use changes in muscular activity and associated shifts in the resonant properties of the bee’s body, to adjust the mechanical properties of their vibrations.

### Function of bee vibrations

In addition to considering differences in the actions of the muscles, another approach to thinking about *why* the muscles produce vibrations with these particular properties is to consider how what properties might best serve these functions. In vibratory communication, for example, the properties of the signalling environment, such as the degree of frequency filtering, determine the “best” vibratory properties to transmit information from producer to receiver (Cocroft & Rodríguez, 2005). Similar factors could influence the “best” properties for defence buzzes. Like the vibratory signals studied in other insect species, the function of a defence buzz is to transmit information from the producer (the bee) to a receiver (the predator). This information is effective; defence or alarm sounds produced by insects, including bumblebees, have been shown to reduce or slow down predator attacks (Masters, 1979; Moore & Hassall, 2016). The effectiveness of defence buzzes is likely affected by the properties of the vibration itself. Although, in our experiment, we found that defence buzzes were on average of lower frequency, peak amplitude velocity and peak amplitude acceleration than floral buzzes, these properties do not correlate with what is likely a more important property of a warning signal: volume (De Luca et al., 2018). A previous comparison of the acoustic properties of defence and floral buzzes found that defence buzzes were significantly louder than floral buzzes (De Luca et al., 2014), and it is possible that the lower frequency or amplitude of the bee’s vibrations during defence buzzing might actually increase the perceived volume of the buzz by predators. A lower frequency and velocity vibration may also be beneficial for the bee as it might be less energetically costly than the higher frequency and velocity floral buzz. Although the costs of buzzing by bees have only been measured for a handful of behaviours (Kammer & Heinrich, 1974; Heinrich, 1975), increasing the frequency and amplitude of vibrations appears to carry a significant cost. For instance, the carpenter bee *Xylocopa varipuncta* Patton, increase the frequency and amplitude of their wingbeats when flying in less dense gases, but doing so increases their metabolic rate by over a third (Roberts et al., 2004). By using lower frequency and velocity vibrations, bumblebees might be able to perform defence buzzes for longer, increasing their effectiveness against predators.

Unlike defence buzzes, the primary function of floral buzzes is not to transmit information to receivers but to shake pollen loose from flowers. Pollen is essential for larval nutrition (Westerkamp, 1996), and bumblebees possess many specialisations to assist in pollen collection, from morphological features such as corbiculae (Thorp, 1979), to behaviour specialisations, including optimising pollen collection (Rasheed & Harder, 1997), rejecting flowers that appear empty of pollen (Buchmann & Cane, 1989; Harder, 1990), and modifying their buzzes in response to the presence or absence of pollen (Russell et al., 2016; Switzer et al., 2019). It is possible that the properties of floral buzzes are also tuned to maximise the pollen collected from poricidal anthers. If that was the case, we would expect the properties that defined floral buzzes in this study, high frequency, velocity, and acceleration, to correlate with the vibration properties which release the most pollen. Studies with artificial shakers have subjected buzz-pollinated flowers to a broad array of vibrations to determine what kinds of vibration release the most pollen (De Luca et al., 2013; Harder & Barclay, 1994; Rosi-Denadai et al., 2018). Although the frequency of floral buzzes appears very consistent across studies, frequency does not appear to determine how much pollen is released from anthers. Instead, as we observed, higher frequencies may result in higher velocities and accelerations, and it is these properties which most determine how much pollen an anther releases (De Luca et al., 2013; Rosi-Denadai et al., 2018). The effect of increasing the velocity or acceleration of floral buzzes on pollen release can be dramatic. De Luca et al. (2013) for example, found that for a floral buzz lasting for one second, doubling the velocity of the buzz led to four times as much pollen being released. Rosi-Denadai et al. (2018) found a similar effect for acceleration – vibrations with a similar acceleration to the floral buzzes we recorded (500 m/s^2^) released more than three times as much pollen as vibrations matching the flight vibrations we recorded (100 m/s^2^), and twice as much as vibrations matching the defence buzzes (300 m/s^2^). The accelerations we recorded from floral buzzes, therefore, are what might be expected from vibrations tuned to maximise pollen release. Producing high acceleration floral buzzes, however, is likely to have come with a cost. Although it is not clear exactly how costly these floral buzzes might be, as no-one has yet measured the metabolic cost of floral buzzing, it has been suggested that bees work to maximise the efficiency of their pollen collection (Rasheed & Harder, 1997). Their foraging decisions are therefore not just based on maximising the pollen their collect, but also based on the potential cost. If floral buzzing exerts a significant cost on bees, this cost might play an important role in their decisions about where and when to forage on buzz-pollinated flowers (Stephens, 2008).

## Conclusion

Our results, demonstrating clear differences between the vibrations by bumblebees in different contexts, open up questions about the mechanisms and evolution of insect vibrations. Currently the mechanisms which control the properties of thoracic vibrations have only been studied in a handful of contexts (Esch & Goller, 1991; King et al., 1996), with most of what we know coming from studies of flight control in *Drosophila* (Lehmann & Bartussek, 2017; Lindsay et al., 2017). The vibrations that individual bumblebees produce in different contexts exhibit stark but reliable differences in their properties, providing a model to better understand how individual insects control the properties of the vibrations they produce. By identifying homologous mechanisms as well as outlining possible constraints on how insect vibrations respond to selection, investigating the mechanisms of bumblebee vibrations can also tell us more about how these behaviours evolve. But to understand how selection might have acted on these vibrations, it is also necessary to examine how bees use these vibrations for their particular functions. The biomechanical properties of a vibration might only be part of what makes it effective. Other behaviours can increase the effectiveness of a particular vibration by increasing the salience or memorability of a signal, such as when animals combine multiple modalities into a signal (Rowe, 1999), or by modifying the effects of the vibrations, such when tree crickets build acoustic baffles to amplify the volume of their mating calls (Mhatre et al., 2017). During floral buzzing, bees do not simply applying vibrations like the artificial shakers used to study pollen release. Instead, bees need to learn to handle flowers correctly, and work to get in position before starting buzzing (Laverty, 1980; Macior, 1964; Russell et al., 2016). How bees handle flowers, where they bite anthers, and how they position themselves as they vibrate, could all influence how the high acceleration vibrations we recorded are applied to the flower and result in pollen ejection. The next step for understanding why bumblebees, and other insects, produce the vibrations they do, is to understand how other behaviours work alongside vibrations to serve their function.

## Acknowledgements

We thank Fernando Montealegre-Zapata and Gema Martin-Ordas for helpful discussions on biomechanics, insects and behaviour. We thank all of the Vallejo-Marín Lab, particularly Carlos Eduardo Pereira-Nunes and Jurene Kemp, for their help with bee maintenance and useful conversations about buzz pollination, and Shoko Sugasawa for comments on the manuscript. This work was supported by a SPARK grant from the University of Stirling and by grant RPG-2018-235 from the Leverhulme Trust.

## Supplementary Materials

**Figure S1:**
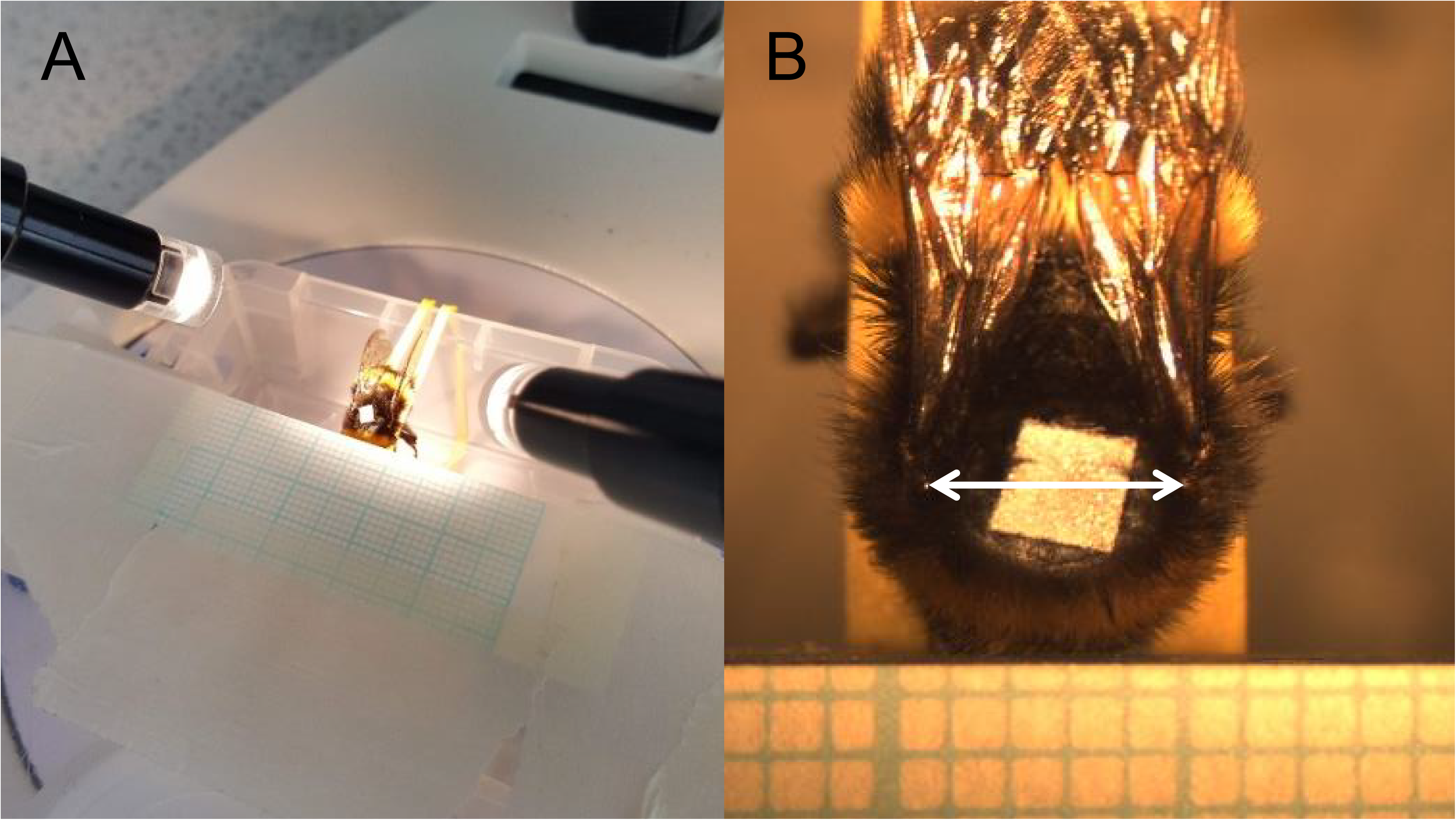
Procedure for assessing bee size. Euthanised bees were held level with graph paper using rubber bands (left), such that a clear image of the intertegular distance (right, white arrow) could be captured by the camera-mounted microscope. The resulting images were then analysed using Image J., using the graph paper to assess the intertegular distance in mm.

**Figure S2.**
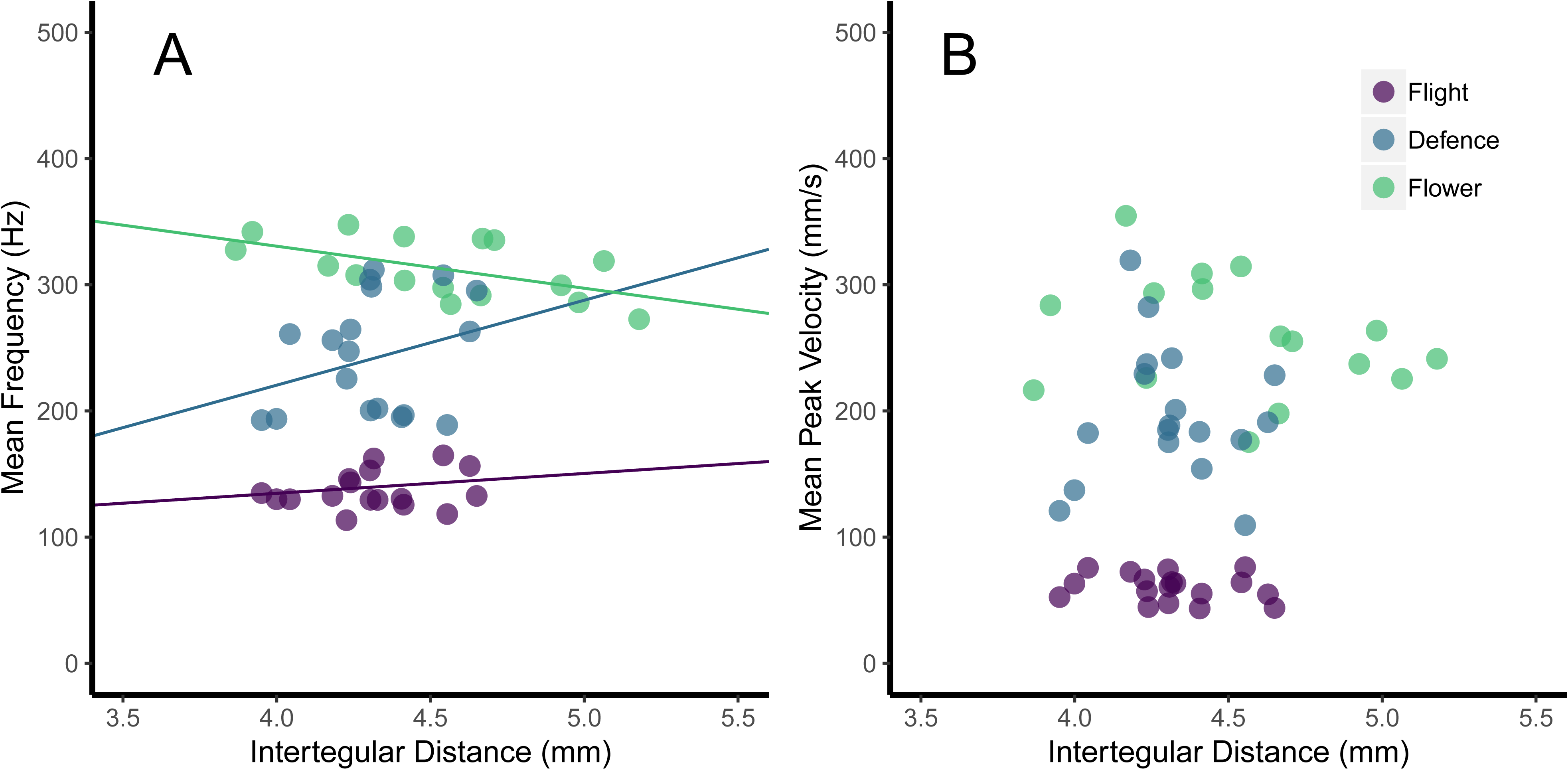
The effect of bee size (in terms of intertegular distance) and behavioural context (flower buzzing, defence and flight) on the frequency (A) and peak velocity (B) of thoracic vibrations. For frequency (A), There was a significant interaction between bee size and behavioural context, with larger bees producing higher frequency defence buzzes, slightly higher frequency flight vibrations, but lower frequency flower buzzes. There was no equivalent interaction for peak velocity (B), with all types of buzzes showing a slight decrease in velocity as bee size increased, although this was not significant (Table 1).

